# Rescue of DNA damage in cells after constricted migration reveals bimodal mechano-regulation of cell cycle

**DOI:** 10.1101/508200

**Authors:** Yuntao Xia, Charlotte R. Pfeifer, Kuangzheng Zhu, Jerome Irianto, Dazhen Liu, Kalia Pannell, Emily J. Chen, Lawrence J. Dooling, Roger A. Greenberg, Dennis E. Discher

## Abstract

Migration through constrictions can clearly rupture nuclei and mis-localize nuclear proteins but damage to DNA remains uncertain as does any effect on cell cycle. Here, myosin-II inhibition rescues rupture and partially rescues the DNA damage marker γH2AX, but an apparent delay in cell cycle is unaffected. Co-overexpression of multiple DNA repair factors and antioxidant inhibition of break formation also have partial effects, independent of rupture. However, there seems to be a bimodal dependence of cell cycle on DNA damage. Migration through custom-etched pores yields the same bimodal, with ~4-μm pores causing intermediate levels of damage and cell cycle delay. Micronuclei (generated in faulty division) of the smallest diameter appear similar to ruptured nuclei, with high DNA damage and entry of chromatin-binding cGAS (cyclic-GMP-AMP-synthase) from cytoplasm but low repair factor levels. Increased genomic variation after constricted migration is quantified in expanding clones and is consistent with (mis)repair of excess DNA damage and subsequent proliferation.

## Introduction

“Go-or-grow” is a hypothesis that posits cell migration and cell cycle are mutually exclusive in space and time (Garay et al., 2013; Giese et al., 1996). For some aspects of 3D invasion, mechanisms of go-or-grow are increasingly being studied with Transwell pores (Beadle et al., 2008; Harada et al., 2014; Wolf et al., 2013). For large pores, migration from contact-inhibited monolayers into sparse microenvironments seems to promote cell cycle re-entry – perhaps analogous to effects of mechanical strain (Benham-Pyle et al., 2015), but for highly constricting pores (Fig.1A), migration appears to delay cell cycle and suppress mitotic counts (Pfeifer et al., 2018). Constricted migration certainly causes nuclear rupture (Denais et al., 2016; Irianto et al., 2017; Raab et al., 2016), and DNA damage seems to increase based on immunostained foci of γH2AX (i.e. phospho-Histone-2AX). However, parallel quantitation of another DNA damage marker (53BP1) shows no increase (Irianto et al., 2017; Pfeifer et al., 2018), with live-cell imaging of large ‘foci’ of GFP-53BP1 (Denais et al., 2016; Raab et al., 2016) attributed to generic accumulation of mobile nuclear protein into low density pockets of chromatin (Irianto et al., 2016). Imaging of DNA damage sites indeed remains non-trivial (Britton et al., 2013), and in the particular context of constricted migration γH2AX foci counts appear to increase by only ~50% across cell cycle stages, even when cell cycle is blocked (Pfeifer et al., 2018). On-the-other-hand, cell cycle checkpoints for DNA damage are well-established (Houtgraaf et al., 2006), and a functional effect of DNA damage on cell cycle in constricted migration could help clarify molecular mechanisms of DNA damage and shed new light on go-or-grow.

**Figure 1.**
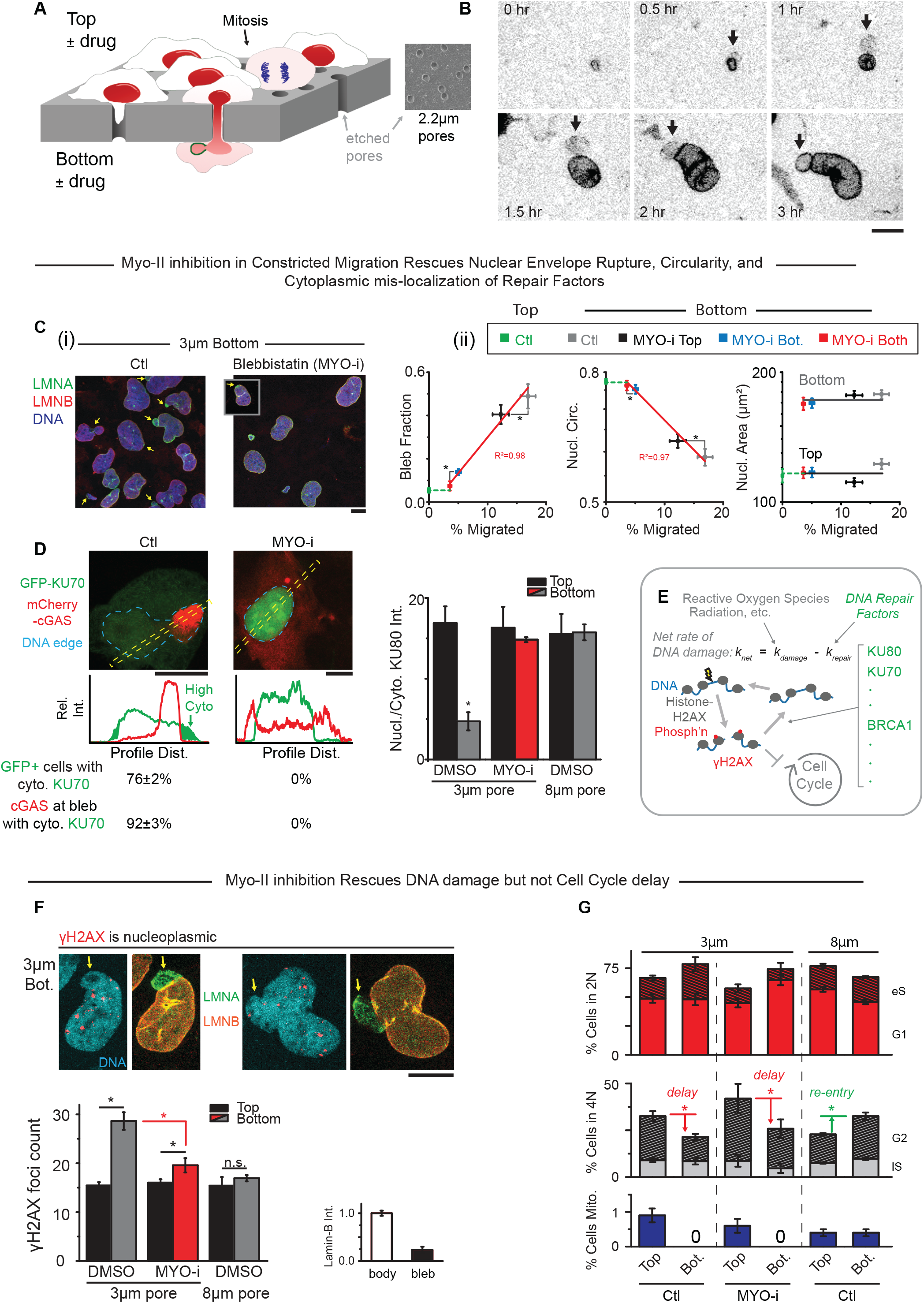
MYO-i on bottom rescues nuclear rupture and DNA damage but not cell cycle delay. (**A**) Nuclei rupture in constricted migration through Transwells that also allow asymmetric exposure to drugs and customized pore size. (**B**) Time-lapse of A549 cell expressing GFP-lamin-A as non-phosphorylatable S22A and emerging from a 3μm pore, with a bleb (arrow) forming at the leading tip of the nucleus. GFP-lamin-A accumulates in the bleb independent of S22 phosphorylation (see also Fig.S1B). (**C**) (**i**) In constricted migration of U2OS cells, nuclear blebs form with lamina disruptions (arrows) except when blebbistatin (MYO-i) is on Bottom. (ii) Addition of MYO-i to the Bottom or Both sides of a Transwell greatly reduces migration and nuclear blebs but increases circularity (*p_xy_<0.05) (>100 cells per condition, n = 5 expt.). (**D**) U2OS cells expressing DNA repair protein GFP-KU70 and DNA-binding protein mCherry-cGAS migrated through 3μm pores; most cells showed GFP-KU70 mis-localized to cytoplasm and mCherry-cGAS accumulated in the nuclear bleb in Ctrl. MYO-i prevents such nucleo-cytoplasmic exchange (n>=2 expt). Bargraph: Endogenous DNA repair factor KU80 also mis-localizes to cytoplasm, except with MYO-i or with larger pores that eliminate blebs (50~300 cells, n>3 expt., *p<0.05). (**E**) DNA breaks constantly form and are repaired, but if net DNA damage is high, then damage checkpoints delay cell cycle. (**F**) Foci of γH2AX (red) are not enriched in nuclear blebs after 3μm pore migration. Bargraphs: γH2AX foci are high except with MYO-i or with larger pores. Compared to the nuclear body, blebs are low in lamin-B as expected. (>100 cells, n=5 expt., *p<0.05). (**G**) Using EdU spike-in to label replicating DNA during Transwell migration, DNA stain intensity and EdU were used to identify a cell as 2N (non-replicated genome) or 4N (fully replicated genome) and as G1, early S (eS), late S (lS), or G2. When contact-inhibited cells migrate through large (8μm) pores into sparse microenvironments, cells re-enter cell cycle. Constricting (3μm) pores delay cell cycle and suppress mitosis - regardless of MYO-i (>400 cells per condition, n=2 expt, *p<0.05). All scalebars: 10μm.

Migration through commercially available constricting pores (3μm diam.) but not large pores (8μm) is driven by myosin-II, with glioblastoma cells upregulating myosin-II and strongly distorting their nucleus while invading through brain slices (Beadle et al., 2008; Ivkovic et al., 2012). Other cell types also seem to use myosin-II to push or perhaps pull the nucleus as a “piston” (Petrie et al., 2014), with inhibition of myosin-II slowing invasion through small circular pores or rectangular channels (Harada et al., 2014; Thiam et al., 2016; Thomas et al., 2015) although activation can also impede invasion (Surcel et al., 2015). Effects of myosin-II on nuclear rupture and DNA integrity remain equally unclear, but actomyosin tension on the front of the nucleus might explain rupture and bleb formation at the front of a squeezed nucleus. Transwells allow drugs to be added to one side or the other to thereby address push or pull mechanisms, and they can also be etched to fine-tune the fit of the nucleus in the pore (Fig.1A).

Nuclear rupture and excess γH2AX has been reported for hyper-contractile cancer cells in 2D culture independent of pore migration (Takaki et al., 2017), and cytoplasmic nucleases have been speculated to enter ‘blebs’ of chromatin at rupture sites during constricted migration (Denais et al., 2016; Raab et al., 2016). Although blebs show no excess γH2AX (Irianto et al., 2017), entry of cGAS (cyclic-GMP-AMP-synthase) into the bleb from cytoplasm might bind γH2AX and inhibit repair (Liu et al., 2018a). DNA repair factors clearly mis-localize to cytoplasm based on immunostaining and live-imaging of GFP constructs, but it remains unknown whether any DNA damage relates to nuclear depletion of such factors or to nuclear entry of other factors. Similar questions apply to micronuclei that form in mitosis with mis-segregated chromosomes (independent of actomyosin) and that somehow exhibit DNA damage while lacking lamin-B as well as multiple DNA repair factors (Hatch et al., 2013; Liu et al., 2018b). We address whether such dysregulation varies with micronuclei diameter in light of pore diameter effects on the main nucleus in migration. Genome variation in micronuclei (Zhang et al., 2015) and in hyper-contractile cancer cells (Takaki et al., 2017) provide evidence of DNA damage through its potential long-term effects, and we examine such effects here after constricted migration. We begin, however, with molecular rescue approaches to a go-or-grow hypothesis that migration through small pores causes myosin-dependent nuclear rupture, repair factor loss, DNA damage, and bimodal delays in cell cycle.

## Results and Discussion

### Myosin-II inhibition rescues nuclear envelope rupture and some DNA damage

As a cell forces its nucleus through a highly constricting pore (3μm diam. for ~10μm length), large distortions last for hours but a nuclear bleb forms early near the highly curved leading edge (Fig. 1B), consistent with past observations of nuclear rupture in migration (Denais et al., 2016; Harada et al., 2014; Irianto et al., 2017; Raab et al., 2016). Lamin-A accumulates slowly on the bleb regardless of phosphorylation state (based on real-time imaging of GFP constructs (Fig.1B) and on immunostaining (Fig.S1A,B)), but such blebs always lack lamin-B (Fig.1C-i) as shown previously. Inhibition of myosin-II with blebbistatin eliminates such blebs while decreasing migration (Fig.1C-ii) and causes nuclei to be more rounded and slightly less spread, consistent with inhibitor effects on many cell types adhering to 2D rigid substrates (Khatau et al., 2009). On such substrates, the MTOC is generally in front of a migrating nucleus and sets the direction for migration (Gomes et al., 2005; Raab and Discher, 2017). Constricting pores, in contrast, set direction and the MTOC typically lags (Fig.S1C). Actin-rich protrusions lead the way in 3D migration and are followed often by myosin-II assembly (Mogilner and Odde, 2011); inhibiting myosin-II only on the bottom of a Transwell is indeed sufficient to suppress nuclear rupture and overall migration (Fig.1C-ii; Fig.S1D-i). Little effect is seen with drug only on top, consistent with minimal diffusion of drug through pores (Fig.S1D-i&ii). Results thus suggest myosin-II helps pull a nucleus through a small pore and ruptures the front of a spreading nucleus (of high area: Fig.1C-ii).

Myosin-II inhibition eliminates mis-localization of nuclear factors into the cytoplasm, such as DNA repair factors KU70 (as GFP construct) and KU80 (endogenous) (Fig.1D, S1E). Mis-localization coincides with entry into the nuclear bleb of cytoplasmic DNA-binding protein mCherry-cGAS (Fig.1D). Large pores (8μm diam.) still require large distortions of nuclei but do not cause mis-localization of nuclear factors (Fig.1D-bargraph). DNA repair factors help maintain DNA damage at low steady state levels, but elevated levels can delay cell cycle (Fig.1E) (Sancar et al., 2004).

Foci of γH2AX in fixed and immunostained nuclei rarely localize to nuclear blebs, that are low in lamin-B, but – more importantly – myosin-II inhibition suppresses total foci counts (Fig.1F-bargraph, S1F). Large pores do not affect a basal level of γH2AX foci, just as they had no effect on rupture (Fig.1D). Nonetheless, a persistent excess of DNA damage even within a non-ruptured, myosin-II inhibited nucleus is consistent with segregation of nuclear factors away from chromatin in constricted migration (Bennett et al., 2017; Irianto et al., 2016). However, functional evidence of DNA damage and its rescue seemed key.

Cell cycle progression, measured as %cells with duplicated genomes (i.e. 4N rather than 2N), increases after cells migrate through 8μm pores from the Transwell top where cells are crowded and contact inhibited (Fig.1G). In contrast, a decrease in %4N is observed after cells migrate through 3μm pores, and no mitotic cells are detected (Fig.1G). Although cell cycle might seem delayed by excess DNA damage, the delay is unaffected by blebbistatin’s *partial* rescue of DNA damage (Fig.1G).

### Overexpressed DNA repair factors or added antioxidant rescues some DNA damage

Mis-localization of DNA repair factors to cytoplasm lasts hours after nuclear envelope rupture, based on imaging of GFP constructs in live cells on rigid substrates (Xia et al., 2018). Nuclear/Cytoplasm intensity ratios for multiple repair factors in cells fixed and immunostained indeed remain low after migration (for 24 hrs) through constricting pores relative to large pores and also relative to cells that have not migrated (Fig.2A,B). A much smaller YFP-NLS construct shows a Nuclear/Cytoplasm intensity that is only slightly low (Fig. 2B), consistent with comparatively rapid nuclear re-entry of such constructs (Raab et al., 2016). We hypothesized that some migration-induced DNA damage could be rescued by overexpression of DNA repair factors because partial knockdown of some DNA repair factors (KU80, BRCA1, etc.) but not others (53BP1) increases DNA damage, and such damage seems additive (Irianto et al., 2017). Indeed, simultaneous co-overexpression of KU70, KU80, and BRCA1 (as GFP constructs) partially rescued the migration-induced DNA damage whereas overexpression of the individual factors as well as GFP-53BP1 showed no significant effects (Fig.2C). Nuclear blebs were still apparent, suggesting that the elevated nuclear levels of multiple key factors facilitated the repair. However, a cell cycle defect in terms of 4N suppression is still seen after 3μm pore migration (Fig. 2D).

**Figure 2.**
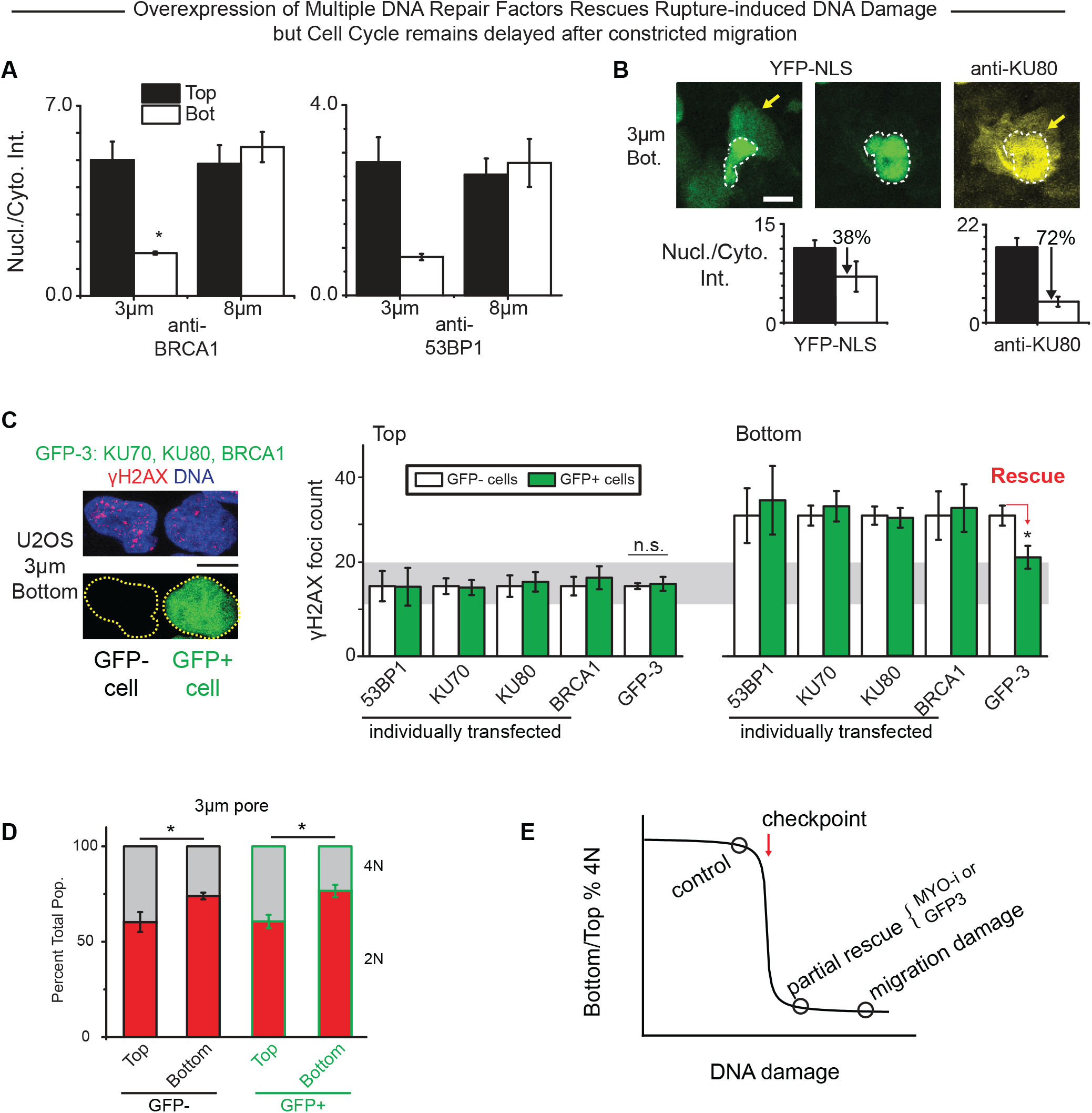
Overexpression of multiple DNA repair factors rescues DNA damage independent of rupture in constricted migration but cell cycle delay persists. (**A**) Constricted migration causes DNA repair proteins BRCA1 and 53BP1 to mis-localize from the nucleus into the cytoplasm (>5 field of views each. n>3 expt., *p<0.05). (**B**) As a nucleus squeezes out of a constriction, both endogenous KU80 (yellow) and overexpressed YFP-NLS (green) are mis-localized to cytoplasm, but nuclei that have fully exited show KU80 remains cytoplasmic while YFP-NLS is mostly nuclear. Bargraphs: Overall Nucl./Cyto. ratio is thus lower for KU80 than for YFP-NLS after 3μm pore migration (>5 field of views each. n>3 expt. per condition, *p<0.05). (**C**) Co-overexpression of three DNA repair factors (GFP-3: KU70, KU80, BRCA1) partially rescues excess DNA damage after constricted migration despite nuclear rupture, whereas rescue is not achieved by overexpression of any single repair protein. (**D**) The 4N population of both Ctl and GFP-3 cells remain suppressed despite the partial rescue of migration-induced DNA damage (200-500 cells per condition. n>3 expt. per condition, *p<0.05). All scalebars: 10 μm. (**E**) A critical amount of DNA damage is predicted to suppress cell cycle progression.

Neither myosin-II inhibition nor repair factor overexpression rescued the cell cycle delay, and so we hypothesized the source of DNA damage could be key to revealing a sigmoidal relationship between cell cycle and DNA damage (Fig.2E). Free radicals and other oxidants can permeate membranes and cause DNA damage (Fig.3A) (Imlay et al., 1988; Kryston et al., 2011; Mahaseth and Kuzminov, 2016; Mello Filho et al., 1984), and even a 0.5-hr exposure to H_2_O_2_ increases γH2AX foci counts and DNA damage in single cell electrophoresis (Fig.S2A), complementing evidence of DNA oxidation (Fig.S2B). Excess repair factors can also reduce H_2_O_2_-induced DNA damage in 2D culture (Fig.S2C). After constricted migration, nuclear oxidative stress is ~30% higher based on labeling by a redox dye (Fig.3A). We therefore added a membrane-permeable antioxidant, glutathione, as a reduced monoethyl ester (GSH-MEE) that reduces H_2_O_2_-induced DNA damage in 2D cultures (Fig.3A, S2A); GSH-MEE has likewise been shown to reduce oxidative stress and genomic instability in pluripotent stem cells in 2D cultures (Skamagki et al., 2017). Partial rescue of DNA damage by GSH-MEE does not affect migrated cell numbers or nuclear blebs (Fig.3B,C). The cell cycle delay persists with antioxidant (Fig.3D), which is again consistent with only a partial rescue of DNA damage.

**Figure 3.**
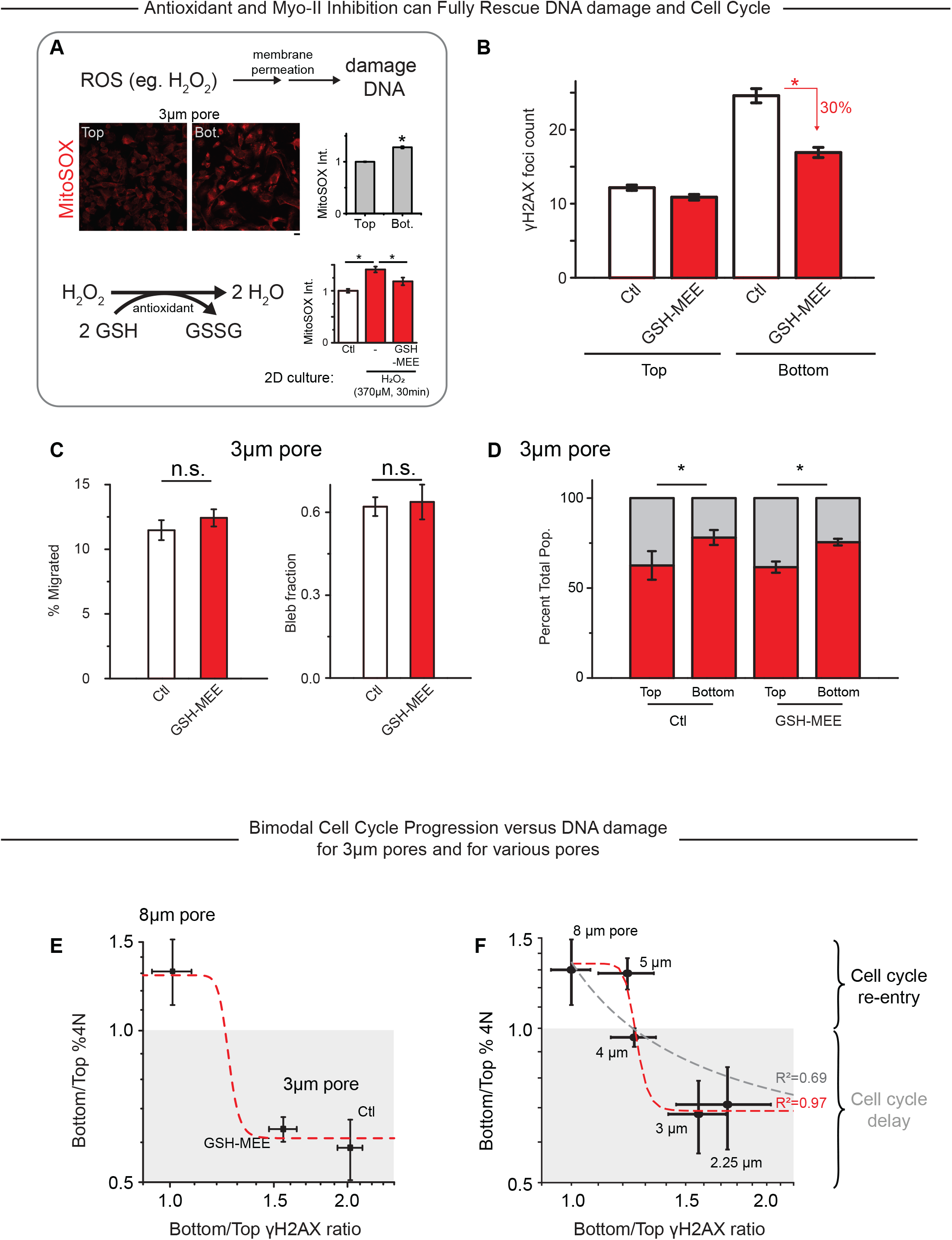
Complete rescue of DNA damage and cell cycle delay requires antioxidant and MYO-i, revealing a bimodal dependence of cell cycle on DNA damage. (**A**) Reactive oxygen species (ROS) damage DNA but can be neutralized by reduced glutathione (GSH), including a membrane-permeable ethyl ester (GSH-MEE). Constricted migration increases ROS based on intensity of MitoSOX dye. (**B**) Cells that treated with such glutathione incur a smaller excess of DNA damage than control cells after constricted migration. (>80 cells per condition, n=3 expt, *p<0.05). All scalebars: l0μm. (**C**) GSH-MEE treatment alone affects neither constricted migration nor nuclear rupture (>3 field of views, n>=2 expt, *p<0.05). (**D**) Both Ctl and GSH-MEE show 4N suppression after constricted migration. (>80 cells per condition, n>=2 expt, *p<0.05). (**E**) Data from panel-D is plotted against the data from panel-B to show %4N remains suppressed on Bottom unless no extra DNA damage is generated. (**F**) For a pore-size dependent test of the cell cycle checkpoint, we enlarged commercial pores by NaOH etching to novel diam’s. of 2.25, 4, and 5μm. Fits use similar *A, B, C* as panel-E with [*m* = 1: *R*^2^ =0.69], [*m* = 3: *R*^2^=0.89], and [*m* = 10: *R*^2^=0.97] (>50 cells per condition, n>=2 expt).

Complete rescue of both cell cycle delay after constricted migration and the DNA damage has not been achieved so far, but a combination of antioxidant with myosin-II inhibition seems promising. As adding the blebbistatin only to the bottom of the Transwell is sufficient to suppress most of the nuclear envelope rupture, the results of DNA damage and cell cycle after adding drugs to Transwell bottom need to be tested as well. However, the results so far suggest that cell cycle progression is a bimodal function of DNA damage caused by constricted migration (Fig. 3E) as proposed in Fig 2E.

### Bimodal cell cycle delay in DNA damage: pore size dependence

To assess the possible generality of a bimodal relationship between cell cycle and DNA damage in migration through pores, we needed to make Transwell pores with a wide range of diameters. Unlike small pores (3μm) studied above, migration through large pores (8μm) promotes cell cycle re-entry and has no effect on basal γH2AX foci counts (Fig.1G) (Pfeifer et al., 2018). Transwells with 2.2μm diameter pores were made from commercially available 1μm pores by adding NaOH to etch the polyester (similarly for 4 & 5μm pores from 3μm pores). After extensive washing, cells are added to the top of the custom-made Transwells as in all studies above, and after 24 hrs, γH2AX foci are counted and cell cycle phases are measured. A pore diameter of 4μm is the critical pore size: relative to cells on top and to large pores, migrated cells show a slight excess in DNA damage and a slight delay in cell cycle, whereas smaller pores (2.25μm, 3μm) cause an increasing excess in DNA damage and a deep but constant delay in cell cycle (Fig.3F). The bimodal results fit a standard Hill equation with a very high cooperativity exponent, *m* >> 1. This typically indicates many interactions upstream (such as in macromolecular γH2AX foci) that amplify signals to downstream regulators.

### Alternative factors: Entry of a nuclease does not affect DNA damage or cell cycle

Other pathways and factors could of course affect DNA damage as well as cell cycle. Entry of cytoplasmic factors into a ruptured nucleus has been speculated to include nucleases (Denais et al., 2016; Raab et al., 2016) and certainly includes cGAS as cited. TREX1 exonuclease and cytoplasmic DNA-binding protein cGAS (*MB21D1* gene) are both expressed in U2OS cells regardless of many rounds of migration (i.e. no selection: Fig.S2D). TREX1 is a transmembrane protein normally in the endoplasmic reticulum (Maciejowski et al., 2015; Rice et al., 2015), and over-expression of functional GFP-TREX1 or an inactive mutant (D18N-TREX1) shows entry into nuclear blebs but no differences in DNA damage on the bleb or in the nucleus before and after constricted migration (Fig.S2E&F); cell cycle delays are also unaffected (Fig.S2G). Endogenous cGAS likewise accumulates in the bleb (Fig.4A; compare also to mCherry-cGAS Fig.1D) and shows no obvious co-localization with γH2AX (Fig.4A, S2H) despite recent suggestions from laser-induced nuclear damage (Liu…Ge Nature 2018). The bleb also immunostains for acetyl-Histone-H3 (Fig.S2I), which is a potential marker of euchromatin (Bannister and Kouzarides, 2011) and which might relate to the restricted binding of cGAS to the bleb.

**Figure 4.**
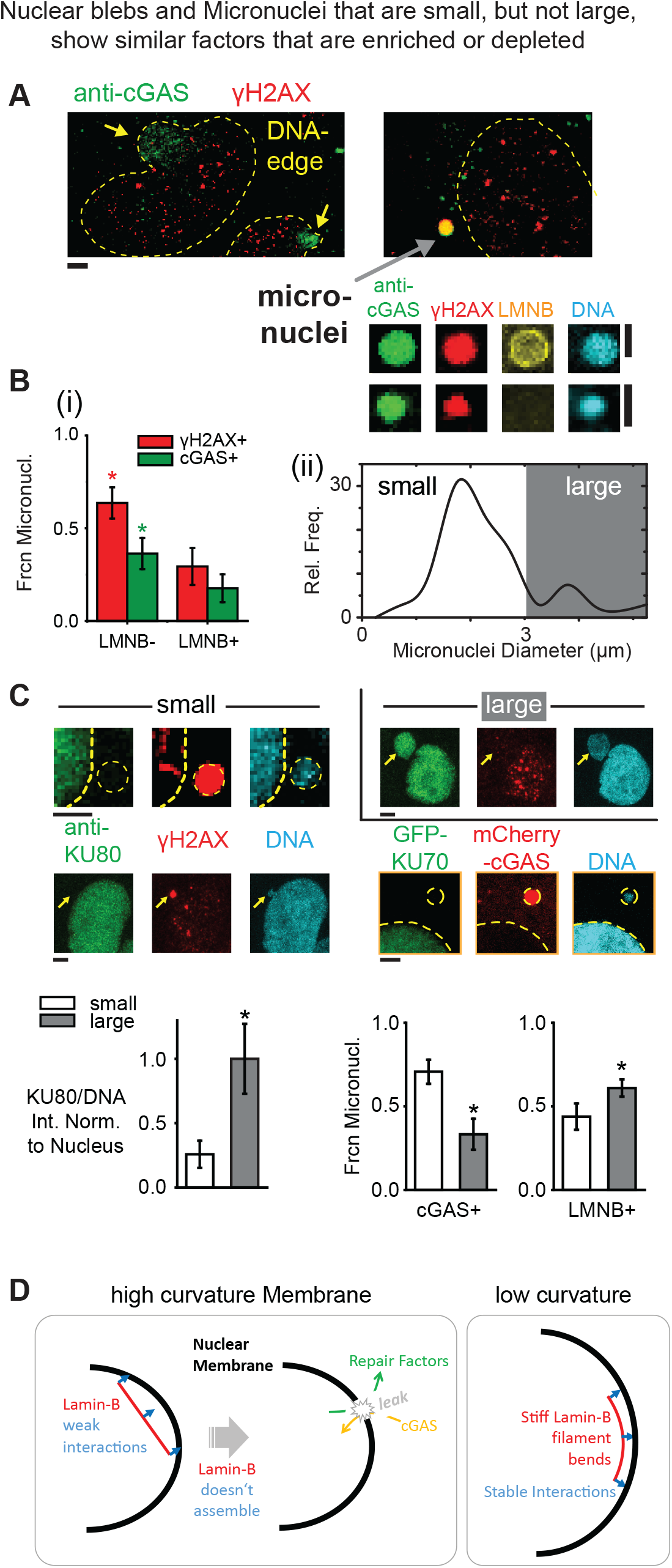
Small micronuclei and ruptured nuclei exhibit many similarities. (**A**) Endogenous cGAS binds DNA in both nuclear blebs (yellow arrows) and many micronuclei (gray arrow). Many micronuclei are γH2AX+ and some lack LMNB. (**B**) Composition of micronuclei assessed after constricted migration shows: (i) More of the low LMNB micronuclei are γH2AX+ and cGAS+. (ii) A broad distribution of small and large micronuclei (n=68). (**C**) Small micronuclei (<3μm) tend to have low KU80 (left 2 images), low GFP-KU70 (bottom right image), low LMNB, but high cGAS (bar graphs). All scalebar: 2μm. (**D**) Lamin-B filaments are too stiff to bend on high curvature nuclear membranes. This applies to small micronuclei and to constricted migration, where blebs form that lack lamin-B.

### Alternative system: Micronuclei of high curvature show less lamin-B and KU80

Nuclear curvature in migration depends of course on pore diameter, and micronuclei generated in cell division also vary in curvature, which potentially provides an alternative approach to studies of nuclear envelope curvature effects. Micronuclei in 2D culture are in particular known to be γH2AX+ and to lack lamin-B, independent of cytoskeleton (Harding et al., 2017; Hatch et al., 2013; Liu et al., 2018b), and DNA-stained micronuclei (Fig.4A) that are most visible within sparsely spread cells on a Transwell bottom tend to be cGAS+ and γH2AX+ when lacking in lamin-B (Fig.4B-i). They also tend to be small (<3μm) though sometimes large (>3μm) (Fig.4B-ii). Small, high curvature micronuclei further lack KU70/KU80 repair factors in addition to being cGAS+ and Lamin-B deficient (Fig.4C).

Importantly, micronuclei can clearly be double-positive as (cGAS+, γH2AX+) (Fig.4A), and so our observations that nuclear blebs are not double-positive (Fig.4A) are unlikely to relate to inhibition of repair by cGAS (Liu et al., 2018a). Another notable difference is that antioxidant fails to affect DNA damage in micronuclei (Fig.S3A). Nonetheless, given the many structural similarities between small micronuclei and nuclear blebs after constricted migration, the fact that lamin-B filaments are stiff (with a persistence length of ~0.5μm or larger (Turgay et al., 2017)) and have high affinity for nuclear envelope (because of farnesylation and lamin-B receptor, LBR) suggests that high curvature membranes energetically oppose stable assembly of lamin-B to thereby favor rupture (Fig.4D,S3B-i&ii).

### Reversible cell cycle delay and Genomic Variation after constricted migration

Delays in cell cycle due to DNA damage could in principle lead to cell senescence or apoptosis, but total cell numbers are nearly constant after 24h of migration, and we do not detect floating cells that commonly indicate cell death. For cells that survive and proliferate after repair of DNA damage, mis-repair can lead to chromosomal abnormalities. This is evident, for example, in sequencing of micronuclei (Liu et al., 2018b) and in mitotic spreads of hypercontractile cancer cells (Takaki et al., 2017). Because U2OS cells are known for high genomic instability and many micronuclei (Zhang et al., 2015), we used A549 lung cancer cells that show in constricted migration similar levels of nuclear envelope rupture, DNA damage, and cell cycle delay (Fig.5A). Cells were migrated or not through small or large pores, detached and expanded (with normal growth rates), migrated again for a total of three times, and then plated as single cells to obtain ~10^6^ cells for genomic analyses (Fig.5B). Expansion in subsequent culture provides some evidence of the reversibility and repair of damage after constricted migration. Chromosome copy number variations (ΔCN) were then quantified using Single Nucleotide Polymorphism (SNP) arrays relative to a technical noise threshold (ΔCN ~ 40 Mb) for duplicate samples and mixtures of cells with different genomes (Fig.S3C). Although all clones obtained after migration through 8 μm pores showed ΔCN within the noise threshold, three of five clones obtained after migration through 3 μm pores showed ΔCN above the noise threshold and up to ~200 Mb (Fig.5C).

**Figure 5.**
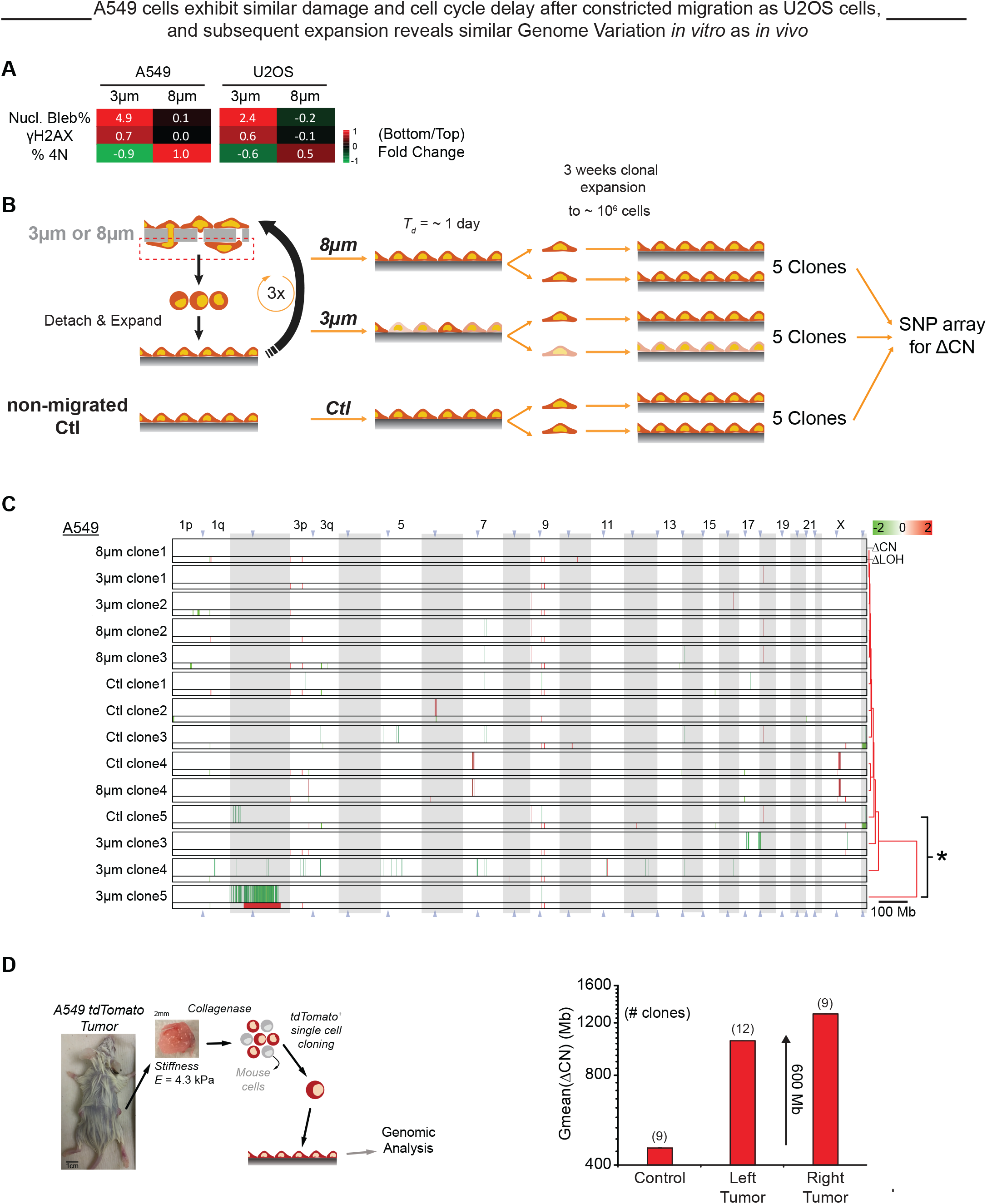
Cell expansion after constricted migration reveals similar genome variation *in vitro* as *in vivo*. (**A**) Heat maps summarize the Bottom-to-Top fold-change in Nucl. Bleb %, DNA damage foci, and % 4N for 3 and 8-μm pore migration of both A549 and U2OS cells (log_2_Ratio). (**B**) Schematics: A549 cells were subjected to three rounds of Transwell migration through 3 or 8μm pores to test the hypothesis that at least some DNA damage would be survivable but mis-repaired. Non-migrated Ctl clones were expanded in parallel. From among these thrice-migrated or non-migrated cells, the genomes of multiple single-cell-derived clones were quantified by SNP array analysis. (**C**) Compared to a clone that migrated three times through 8μm pores, significant chromosome copy number changes (ΔCN) and loss of heterozygosity (ΔLOH) above the noise level (40 Mb) are observed in 3 out of 5 A549 clones that migrated through 3 μm pores. Clones are listed per hierarchical clustering (city-block metric) of their ΔCN, and the asterisk indicates statistical significance in the overall distribution of gains (red) and losses (green) (*p<0.05 in Kolmogorov-Smirnov test). (**D**) Schematic: A549 tdTomato cells were injected subcutaneously into the left and right flanks of non-obese diabetic/severe combined immunodeficient (NOD/SCID) mice (n⩾3 mice). After 4 weeks, tumors were harvested and disaggregated, and SNP array analysis was performed on single-cell-derived tdTomato^+^ clones. Bargraph: Clones from both the left and right tumors exhibited significant ΔCN compared to control cells cultured on 2D plastic. This increase in genomic variation is similar in magnitude to that observed after constricted migration.

While *in vitro* studies allowed tight control over many variables, *in vivo* variation provides an important benchmark: ΔCN might prove very low, for example. Meta-analyses of complex human tumors further indicate that tumor cells in stiff tissues tend to harbor more mutations than tumors in soft tissues (Pfeifer et al., 2017), and stiffer tissues have more collagen and should therefore have lower porosity. Tumors of lung cancer-derived A549 cells in immunodeficient mice (NSG) do not interact with cells of the acquired immune system that limit growth of lung tumors with high genomic variation (Rizvi et al., 2015). Compared to control cultures that proliferated for similar duration but never underwent 3D migration, A549 tumors in stiff subcutaneous sites exhibit greater genomic variation (by ΔCN ~ 600 Mb) (Fig. 5D), which is similar in magnitude as that achievable in more controlled *in vitro* studies of constricted migration.

## Conclusions

Nuclear rupture has been clearly demonstrated during cell migration *in vivo* with YFP-NLS, and findings here with rigid Transwells suggest similar studies should provide further insight when using GFP constructs of repair factors (Fig.2A) and cell cycle markers. Transwells require the cell and nucleus to deform strongly, but deformable channels (Pathak and Kumar, 2012) might reveal a smaller equivalent diameter than the critical diameter reported here (Fig.3F). Transwells are also impermeable to liquid flow, and so roles for aquaporins in migration through impermeable channels are also likely (Stroka et al., 2014) and would also be important to compare to roles *in vivo*. Effects of myosin-II in constricted migration seem generally consistent with results for hypercontractile cells in 2D rigid cultures (Takaki et al., 2017), which also suggested genome variation even though myosin-II knockout in yeast (a stationary cell type) has the opposite effect in increasing aneuploidy (Rancati et al., 2008). High curvature regions of interphase nuclei in 2D culture are prone to rupture and DNA damage, which are favored by high contractility and by rigid substrates but independent of migration (Xia et al., 2018), consistent with effects of both constricting pores and micronuclei of high curvature. Lamin-B filaments are stiff, with a persistence length of ~0.5μm (Turgay et al., 2017), and therefore will not bend and bind a nuclear envelope with high Gaussian curvature, but myosin tension is somehow independently important because pores impose constant curvature. Squeezing of dozens of mobile proteins into low density regions of chromatin could also be relevant and likely explains the large intranuclear accumulations of GFP-53BP1 during migration through narrow channels (Denais et al., 2016; Irianto et al., 2017; Raab et al., 2016) independent of DNA damage (Irianto et al., 2017; Pfeifer et al., 2018). DNA damage is nonetheless consistent with simple orthogonal measures of a bimodal cell cycle delay, illustrating mechanical control of cell cycle as one mechanism of “go-or-grow”.

## Materials and Methods

### Cell culture

U2OS human osteosarcoma cells and A549 human lung carcinoma cells were cultured in DMEM high-glucose medium and Ham’s F12 medium (Gibco, Life Technologies), respectively, supplemented with 10% FBS and 1% penicillin/streptomycin (MilliporeSigma). Cells were incubated at 37°C and 5% CO_2_, as recommended by ATCC.

### Immunostaining

Cells were fixed in 4% formaldehyde (MilliporeSigma) for 15 minutes, followed by 15-minute permeabilization by 0.5% Triton-X (MilliporeSigma), 30-minute blocking by 5% BSA (MilliporeSigma), and overnight incubation in primary antibodies at 4°C. The antibodies used include lamin-A/C (1:500, sc-7292, mouse, Santa Cruz), lamin-B (1:500, goat, sc-6217, Santa Cruz & 1:500, rabbit, ab16048, Abcam), γH2AX (1:500, mouse, 05-636-I, MilliporeSigma), 53BP1 (1:300, rabbit, NB100-304, Novus), KU80 (1:500, rabbit, C48E7, Cell Signaling), BRCA1 (1:500, mouse, sc-6954, Santa Cruz), cGas (1:500, rabbit, D1D3G, Cell Signaling), Pericentrin (1:500, rabbit, ab4448, Abcam), Acetyl-H3 (1:500, rabbit, ab47915, ABcam) and RPA2 (1:500, mouse, ab2175, Abcam). Finally, after 90-minute incubation in secondary antibodies (1:500, donkey anti mouse, goat or rabbit, ThermoFisher), the cells’ nuclei were stained with 8 μM Hoechst 33342 (ThermoFisher) for 15 minutes. When used, 1 μg/mL phalloidin-TRITC (MilliporeSigma) was added to cells for 45 minutes just prior to Hoechst staining. MitoSOX is applied to live samples for 10-15 minutes at 3.8*10^-3^ μg/μl before fix. CFDA (Vybrant™ CFDA SE Cell Tracer Kit, Thermo, Cat# V12883) is applied to live cells at 2μM in live samples as instructed.

### EdU labeling and staining

EdU (10 μM, Abcam) was added to Transwell membrane 1 hour before fixation and permeabilization. After permeabilization, samples were stained with 100 mM Tris (pH 8.5) (MilliporeSigma), 1 mM CuSO4 (MilliporeSigma), 100 μM Cy5 azide dye (Cyandye), and 100 mM ascorbic acid (MilliporeSigma) for 30 min at room temperature. Samples were thoroughly washed, and then underwent immunostaining as described above.

### Transwell migration

Migration assays were performed using 24-well inserts with 3 μm and 8 μm-pore filters (Corning) with 2×10^6^ and 1 ×10^5^ pores per cm^2^ respectively. A total of 1.5×10^5^ cells were seeded on top of the pore filters. The media, supplemented with 10% FBS and 1% penicillin-streptomycin, was added to both the top and bottom of each 24-well insert such that there was no nutrient gradient across the pore filter. After incubating for approximately 24 hours at 37°C and 5% CO_2_, the entire filter was fixed and stained with desired antibodies following standard immunofluorescence protocol.

### Transwell pore etching

Porous membrane cell culture inserts are commercially available from Corning; they come in a limited assortment of pore diameters, including 1, 3, 5 and 8 μm. To generate intermediate pore sizes, such as 4 or 6 μm, the polycarbonate membranes were etched with 2 M NaOH in a 60°C incubator. 3 μm pore membranes were incubated for 72 min and 120 min to generate 4 μm and 5 μm diameter pores, respectively. The same conditions were used to etch 5 μm pore membranes to generate 6 and 7 μm diameter pores. To achieve pores of sub-3 μm diameter, 1 μm pore membranes were irradiated with 365 nm ultraviolet light (Spectroline XX-15A, 0.7 A) for 30 min per side, and then etched with 9 M NaOH at room temperature (22°C). Etching for 4.5 hr and 5 hr yielded 2.2 μm and 2.3 μm diameter pores, respectively. In every case, after etching, membranes were thoroughly washed with MilliQ water and dried under vacuum. Etched membranes were sterilized with UV irradiation before being used in migration assays. Etching conditions were adapted from (Cornelius et al., 2007).

### Imaging

Conventional epifluorescence images were taken using an Olympus IX71 microscope—with a 40x/0.6 NA objective—and a digital EMCCD camera (Cascade 512B, Photometrics). Confocal imaging was done on a Leica TCS SP8 system with a 63x/1.4 NA oil-immersion objective. Scanning electron microscopy (SEM) of etched Transwell membranes was performed with an FEI Quanta 600 FEG Mark II ESEM, operated in wet environmental mode.

### Transfection in U2OS cells

GFP-TRX1 WT and D18N are from Addgene (#27219, 27220). GFP-BRCA1 (Addgene plasmid # 71116) was a gift from Dr. Daniel Durocher of the Lunenfeld-Tanenbaum Research Institute in Toronto, Canada; GFP-KU70 and GFP-KU80 were gifts from Dr. Stuart L. Rulten of the University of Sussex in Brighton, UK (Grundy et al., 2013); and mCherry-cGAS were gifts from Dr. Roger Greenberg (Harding et al., 2017). GFP-LMNA-S22A was used in our prior studies (Buxboim et al., 2014). Cells were passaged 24 hours prior to transfection. A complex GFPs (0.2-0.5 ng/mL) and 1 μg/mL Lipofectamine 2000 (Invitrogen, Life Technologies) was prepared according to manufacturer instructions, and then added for 24 hours (GFPs) to cells in corresponding media supplemented with 10% FBS. GFP-3 consists of GFP-KU70, GFP-KU80, and GFP-BRCA1 (0.2-0.5 ng/mL each).

### Establishment of A549 Tumors *In Vivo*

For each injection, ~10^6^ cells were suspended in 100 μL ice-cold PBS and 25% Matrigel (BD) and injected subcutaneously into the flank of non-obese diabetic/severe combined immunodeficient (NOD/SCID) mice with null expression of interleukin-2 receptor gamma chain (NSG mice). Mice were obtained from the University of Pennsylvania Stem Cell and Xenograft Core. All animal experiments were planned and performed according to IACUC protocols. The tumors were grown for 4 weeks and tumors cells from mice were then isolated for SNP array.

### Genome (SNP array) analysis

DNA isolation used the Blood & Cell Culture DNA Mini Kit (Qiagen) per manufacturer’s instruction. The same DNA samples were also sent to The Center for Applied Genomics Core in The Children’s Hospital of Philadelphia, PA, for Single Nucleotide Polymorphism (SNP) array HumanOmniExpress-24 BeadChip Kit (Illumina). For this array, >700,000 probes have an average inter-probe distance of ~4kb along the entire genome. For each sample, the Genomics Core provided the data in the form of GenomeStudio files (Illumina). Chromosome copy number and LOH regions were analyzed in GenomeStudio by using cnvPartition plug-in (Illumina). Regions with one chromosome copy number are not associated with LOH by the Illumina’s algorithm. Hence, regions with one chromosome copy number as given by the GenomeStudio are added to the LOH region lists. SNP array experiments also provide genotype data, which was used to give Single Nucleotide Variation (SNV) data. In order to increase the confidence of LOH data given by the GenomeStudio, the changes in LOH of each chromosome from each sample were cross referenced to their corresponding SNV data. Comparison analyses between SNPa and aCGH were again done in MATLAB, and in order to compare different samples, probes with ‘no-call’ (either due to low read intensity or located outside the ‘call’ cluster) were removed from further analysis.

### Statistical analyses

All statistical analyses were performed using Microsoft Excel 2013. Unless otherwise noted, statistical comparisons were made by unpaired two-tailed Student *t*-test and were considered significant if *p* < 0.05. *p_xy_* in Fig. 1C-ii is the joint probability obtained by multiplying *p_x_* and *p_y_*, which assumes independence (Wang, 2013). Unless mentioned, all plots show mean ± S.E.M. “n” indicates the number of samples, cells, or wells quantified in each experiment. Figure legends specify the exact meaning of “n” for each figure (Cumming et al., 2007).

## Acknowledgments

The authors would like to thank Dr. Stuart L Rulten from University of Sussex, Brighton, UK for various DNA damage response protein plasmids used in this study and Dr. Qinnan Zhang from University of Notre Dame for schematic drawing. The authors in this study were supported by the National Institutes of Health National Cancer Institute under Physical Sciences Oncology Center Award U54 CA193417, National Heart Lung and Blood Institute Award R21 HL128187, the US–Israel Binational Science Foundation, Charles Kaufman Foundation Award KA2015-79197, and National Science Foundation grant agreement CMMI 15-48571. The content is solely the responsibility of the authors and does not necessarily represent the official views of the National Institutes of Health nor the National Science Foundation. The authors declare no competing financial interests.

## Supplement Figure Legends

**Figure S1.**
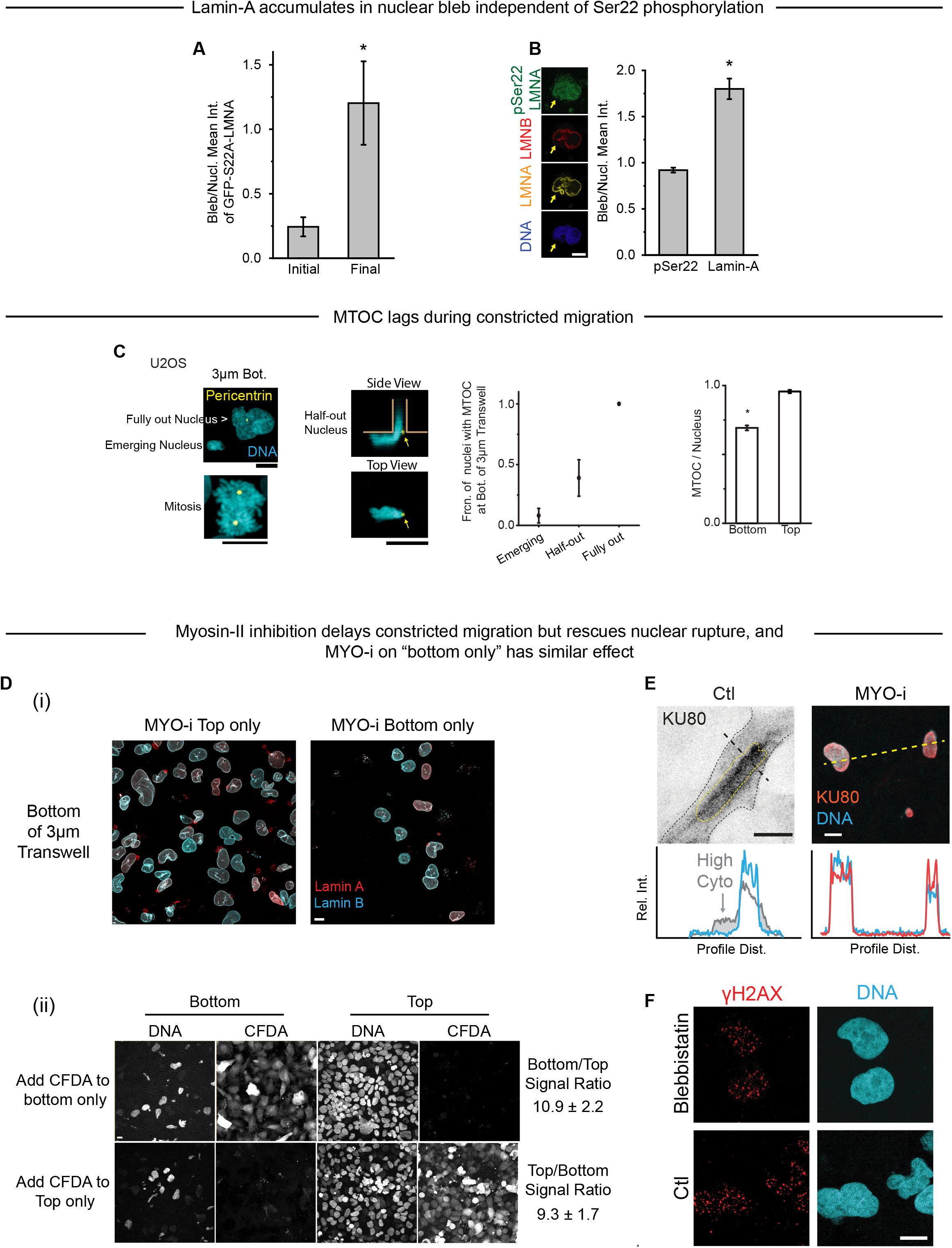
MYO-i rescues nuclear rupture and DNA damage. (**A**) GFP-S22A-LMNA accumulates in nuclear blebs over time, causing the bleb-to-nucleus GFP intensity ratio to increase after bleb formation (n=3 cells, *p<0.05). (**B**) Immunostaining of nuclei on the Bottom of a 3μm pore filter indicates enrichment of lamin-A but not lamin-B or pSer22-lamin-A in nuclear blebs (n>5 cells, *p<0.05). (**C**) MTOC staining (pericentrin) is confirmed to be clean, as MTOCs can be clearly seen in mitotic cells. Some nuclei on the Bottom of a 3μm pore filter do not have visible MTOCs (lower image, yellow arrow), giving Bottom a lower MTOC-per-nucleus rate than Top (bar graph). This lower rate occurs because the MTOC is not typically oriented towards the leading edge of a pore-migrating U2OS nucleus. Hence, in nuclei that are just emerging from pores, MTOCs are not visible on the membrane Bottom. As nuclei emerge further, some MTOCs become visible outside the pore (images of “Half-out Nucleus”). Once the whole nucleus has completely migrated, the MTOC is invariably seen on Bottom (right plot, >48 cells per condition, n>=3 expt, *p<0.05). (**D**) (i) Representative images show U2OS nuclei after migration through 3μm pores. Blebbistatin (MYO-i) was added either only on Top of the 3μm pore filter (left) or only on Bottom (right). (ii) When added only to the Top or Bottom of a Transwell pore filter, small molecules such as the cell-permeable dye CFDA do not severely cross to the other side of the filter (n=3 expt). (**E**) A representative immunofluorescence image shows a ruptured control (Ctl) nucleus with mis-localization of endogenous KU80 from the nucleus into the cytoplasm. After MYO-i treatment, rupture is rescued, as KU80 does not mis-localize. (**F**) Representative images show DNA and DNA damage in 3μm pore-migrated cells with and without MYO-i treatment. All scale bars are 10 μm.

**Figure S2.**
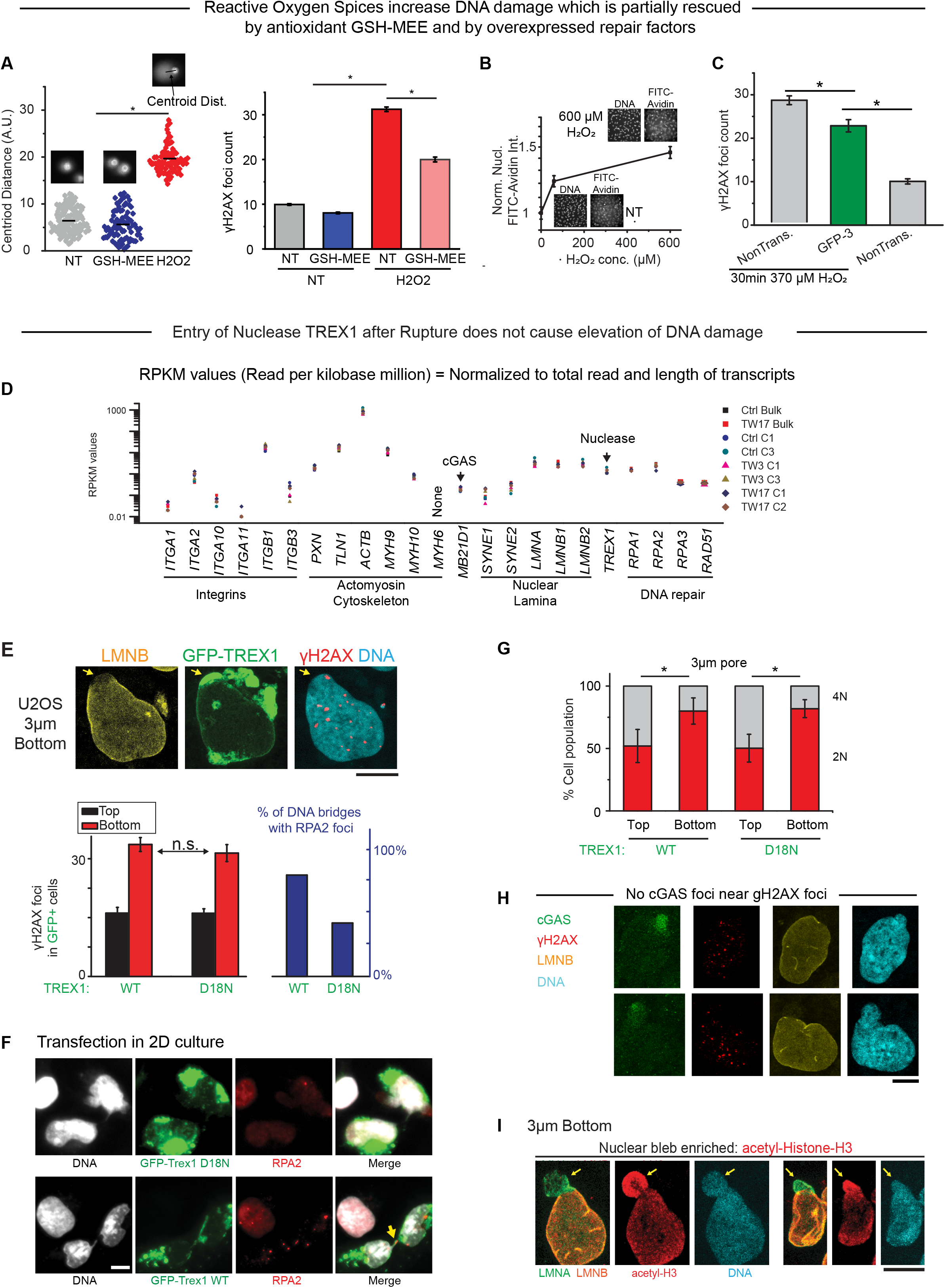
ROS level correlates with DNA damage level. TREX1 entry after rupture does not cause elevation of DNA damage. (**A**) H_2_O_2_ (370μM, 30min) severely damages DNA, creating massive DNA fragments detected by electrophoretic comet assays: nuclei isolated after H_2_O_2_ treatment show more of a cathode-shifted centroid of DNA with a higher mean displacement (inset images) as compared to nuclei isolated from NT and GSH-MEE groups (50-100 cells per condition, n=3 expt, *p<0.05). H_2_O_2_-induced DNA damage is also confirmed by γH2AX foci count. The excess damage can be partially rescued by adding 1mg/mL GSH-MEE (200-700 cells per group, n>3 expt, * p<0.05). (**B**) FITC-labeled avidin has a high specific affinity for 8-oxoguanine, an oxidized base in DNA. Half-hour incubation with 60 or 600 μM H_2_O_2_ results in elevated nuclear FITC-avidin signal, suggesting increased oxidative stress (>120 cells per group, *p<0.05, n=2 expt). (**C**) Co-overexpression of KU70, KU80 and BRCA1 (GFP-3) reduces H_2_O_2_-induced DNA damage, though not to the basal level measured in nontransfected (NonTrans.) cells without H_2_O_2_ (28-132 cells per condition, n= expt, *p<0.05). (**D**) In our previous study (Irianto, et al., 2017), 3μm pore migration of U2OS cells for three (TW3) or seventeen (TW17) rounds was followed by clonal expansion and RNA-Seq analysis. Non-migrated Ctrl clones and bulk cells were expanded in parallel. Here, we find that multiple mechanosensing components remain unperturbed by repeated constricted migration. The RNA-Seq data also reveals expression of at least two cytoplasmic factors, TREX1 and cGAS (TREX1 and MB21D1), that might interact with chromatin exposed by nuclear envelope rupture. (**E**) Overexpressed nuclease GFP-TREX1 localizes to site of nuclear rupture after constricted migration (arrow), but γH2AX foci are not enriched. Bargraphs (left): Both wild-type (WT) and inactive (D18N) GFP-TREX1 show the same excess DNA damage after constricted migration. Functionality of WT TREX1 was confirmed by an increase in RPA2 foci on mitotic bridges of DNA (right) (>150 cells per condition, n = 3 expt., *p<0.05). (**F**) Representative images: Overexpression of wild-type TREX1 (GFP-TREX1-WT) causes formation of RPA2 foci on DNA bridges (yellow arrow) post-mitosis, confirming nuclease activity. With overexpression of inactive mutant TREX1 (GFP-TREX1-D18N), fewer cells exhibit such RPA2 foci. (**G**) Suppression of 4N is the same for WT and mutant GFP-TREX1 (>150 cells per condition, n = 3 expt., *p<0.05). (**H**) Endogenous cGAS binds to DNA in nuclear blebs, as seen in representative images of U2OS cells after 3μm pore migration. DNA damage foci do not localize to blebs, the sites of cGAS accumulation. (**I**) Representative images of 3μm pore-migrated U2OS cells show that nuclear blebs have abundant acetylated chromatin. All scale bars are 10 μm.

**Figure S3.**
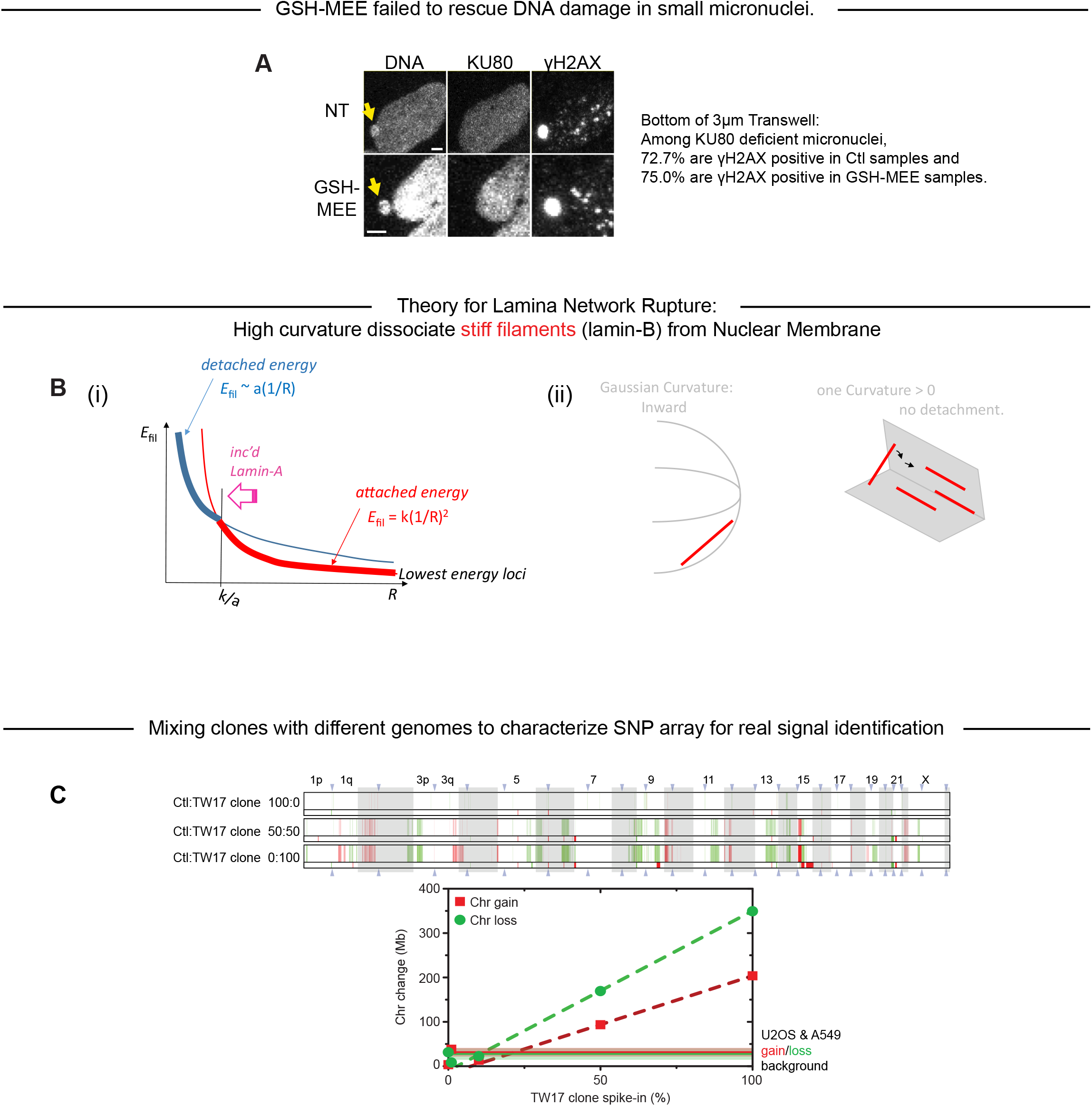
Theory for lamina rupture and characterization of SNP array. (**A**) GSH-MEE does not reduce the fraction of γH2AX-positive micronuclei that lack KU80 (n=10). scale bar is 2 μm. (**B**) (i) When attached to the nuclear membrane, a stiff lamin-B filament of a given length has a bending energy (red); when detached, the same filament has an association energy with the membrane (blue). At large radius of membrane curvature *R*, it is energetically favorable for a lamin-B filament to remain attached to the nuclear membrane (think red curve); at small *R*, it becomes favorable for lamin-B to dissociate (thick blue curve). The attached-to-detached transition occurs at *k*/*a*, where *k* is the bending modulus of an attached filament and *a* is the interaction energy between a detached filament and the nuclear membrane. In each regime (<*k*/*a* and >*k*/*a*), the lower-energy—hence, favorable—state is indicated by a thick curve. (ii) With membranes of high Gaussian curvature, stiff filaments (lamin-B) cannot perfectly conform or cling to the surface, which drives detachment of the filaments. However, for channel-shaped membranes and pores that are flat in on direction, stiff straight filaments can rotate and still cling to the surface, hence no detachment. (**C**) To characterize the signal and noise levels of our SNP array data, two distinguishable U2OS samples—generated in our previous study (Irianto, et al., 2017) —were mixed and subjected to SNPa analysis. The real signal for genomic variation becomes detectable at Chr change > 40 Mb.

